# High-throughput detection of antibodies targeting the SARS-CoV-2 Spike in longitudinal convalescent plasma samples

**DOI:** 10.1101/2020.10.20.346783

**Authors:** Sai Priya Anand, Jérémie Prévost, Jonathan Richard, Josée Perreault, Tony Tremblay, Mathieu Drouin, Marie-Josée Fournier, Antoine Lewin, Renée Bazin, Andrés Finzi

**Affiliations:** Centre de Recherche du CHUM, Montréal, QC, Canada; Department of Microbiology and Immunology, McGill University, Montréal, QC, Canada; Département de Microbiologie, Infectiologie et Immunologie, Université de Montréal, Montréal, QC, Canada; Héma-Québec, Affaires Médicales et Innovation, Québec City / Montréal, QC, Canada

## Abstract

**Background:** The SARS-CoV-2 virus is the cause of the ongoing coronavirus disease 2019 (COVID-19) pandemic, infecting millions of people and causing more than a million deaths. The SARS-CoV-2 Spike glycoproteins mediate viral entry and represent the main target for antibody responses. Humoral responses were shown to be important for preventing and controlling infection by coronaviruses. A promising approach to reduce the severity of COVID-19 is the transfusion of convalescent plasma. However, longitudinal studies revealed that the level of antibodies targeting the receptor-binding domain (RBD) of the SARS-CoV-2 Spike declines rapidly after the resolution of the infection.

**Study Design and Methods:** To extend this observation beyond the RBD domain, we performed a longitudinal analysis of the persistence of antibodies targeting the full-length SARS-CoV-2 Spike in the plasma from 15 convalescent donors. We generated a 293T cell line constitutively expressing the SARS-CoV-2 Spike and used it to develop a high-throughput flow cytometry-based assay to detect SARS-CoV-2 Spike specific antibodies in the plasma of convalescent donors.

**Results and Conclusion:** We found that the level of antibodies targeting the full-length SARS-CoV-2 Spike declines gradually after the resolution of the infection. This decline was not related to the number of donations, but strongly correlated with the decline of RBD-specific antibodies and the number of days post-symptom onset. These findings help to better understand the decline of humoral responses against the SARS-CoV-2 Spike and provide important information on when to collect plasma after recovery from active infection for convalescent plasma transfusion.

## Introduction

The ongoing coronavirus disease 2019 (COVID-19) pandemic is caused by the severe acute respiratory syndrome coronavirus 2 (SARS-CoV-2) and as of October 2020, has caused over a million deaths worldwide (https://www.worldometers.info/coronavirus/). The transfusion of convalescent plasma for the treatment of respiratory infections caused by coronaviruses, such as SARS-CoV-1, has been successful to improve patient outcome ^1^. Its use has now been initiated as an adjunctive therapy for patients with COVID-19 and several clinical trials are underway (for example NCT04412486 and NCT04342182). Preliminary findings have suggested improvements in the patients’ clinical status after convalescent plasma treatment ^2–5^.

Currently, the dynamics of the humoral response against SARS-CoV-2 are under investigation. Of importance is the highly immunogenic trimeric Spike (S) glycoprotein, which is the target of neutralizing antibodies (Abs) and facilitates SARS-CoV-2 entry into host cells via its receptor-binding domain (RBD) that interacts with angiotensin-converting enzyme 2 (ACE-2) ^6,7^. The neutralization activity of plasma from convalescent donors has been suggested to be important for clinical improvement and is a factor of consideration in screening convalescent plasma ^2,3,8,9^. However, several studies have shown that antibody titers and neutralization activity against S, including RBD-specific Abs, decrease during the first weeks after resolution of infection ^10–12^. Furthermore, despite most neutralizing Abs being RBD-specific ^12–14^, studies have isolated potent neutralizing Abs that are specific to other epitopes on the S trimer, mainly directed against the N-terminal domain of the S1 subunit (NTD) ^15^. Additionally, the bulk of the antibody responses elicited by SARS-CoV-2 infection were found to target two major immunodominant regions on the S protein, such as the fusion peptide region and heptad repeat 2 (HR2) of the S2 subunit ^16,17^. Thus, current plasma screening processes using only recombinant RBD to determine seropositivity and antibody titers for convalescent plasma therapy could overlook antibodies specific to multiple epitopes on the viral spike. Here we have developed a high-throughput flow-cytometry assay that is based on the recognition of the full-length SARS-CoV-2 S protein expressed on the surface of 293T cells. This method allows for the detection of antibodies binding to various conformations and domains of the Spike. We used this method to screen longitudinal convalescent plasma samples from 15 donors to determine the antibody response to the full Spike over time.

## Material and Methods

### Convalescent plasma donors

Recovered COVID-19 patients were recruited mostly following self-identification and through social media. All participants have received a diagnosis of COVID-19 by the Québec Provincial Health Authority and met the donor selection criteria for plasma donation in use at Héma-Québec. They donated plasma at least 14 days after complete resolution of COVID-19 symptoms. Males and females with no history of pregnancy meeting the above criteria were invited to donate plasma, after informed consent. A volume of 500 mL to 750 mL of plasma was collected by plasmapheresis (TRIMA Accel®, Terumo BCT). Seropositive donors donated additional plasma units every six days, for a maximum of 12 weeks. All work was conducted in accordance with the Declaration of Helsinki in terms of informed consent and approval by an appropriate institutional board. Convalescent plasmas were obtained from donors who consented to participate in this research project at Héma-Québec (REB # 2020-004).

### Transfection and transduction of 293T cells

293T human embryonic kidney cells (obtained from ATCC) were maintained at 37°C under 5% CO2 in Dulbecco’s modified Eagle’s medium (DMEM) (Wisent) containing 5% fetal bovine serum (VWR) and 100 μg/ml of penicillin-streptomycin (Wisent). The plasmid expressing the full-length SARS-CoV-2 Spike was kindly provided by Stefan Pöhlmann and was previously reported ^7^. 293T cells were transfected with 10 μg of Spike expressor and 2 μg of a green fluorescent protein (GFP) expressor (pIRES-GFP) for 2×10^6^ 293T cells using the standard calcium phosphate method. For the generation of 293T cells stably expressing the SARS-CoV-2 Spike protein, transgenic lentiviruses were produced in 293T using a third-generation lentiviral vector system. Briefly, 293T cells were co-transfected with two packaging plasmids (pLP1 and pLP2), an envelope plasmid (pSVCMV-IN-VSV-G) and a lentiviral transfer plasmid coding for a GFP-tagged SARS-CoV-2 Spike (pLV-SARS-CoV-2 S C-GFPSpark tag) (Sino Biological). Supernatant containing lentiviral particles was used to transduce more 293T cells in presence of 5μg/mL polybrene. The 293T cells stably expressing SARS-CoV-2 Spike (GFP+) were sorted by flow cytometry.

### Cell surface staining and flow cytometry analysis

293T cells transfected with a Spike expressor or 293T-Spike cells were stained with the anti-RBD CR3022 monoclonal Ab (5 μg/ml) or plasma (1:250 dilution). AlexaFluor-647-conjugated goat anti-human IgG (H+L) Abs (Invitrogen) were used as secondary antibodies. The percentage of transfected/transduced cells (GFP+ cells) was determined by gating the living cell population based on viability dye staining (Aqua Vivid, Invitrogen). Samples were acquired on a LSRII cytometer (BD Biosciences) and data analysis was performed using FlowJo v10.5.3 (Tree Star). The seropositivity threshold was established using the following formula: (mean of all COVID-19 negative plasma + (3 standard deviation of the mean of all COVID-19 negative plasma) + inter-assay coefficient of variability).

### Statistical analyses

Statistics were analyzed using GraphPad Prism version 8.4.3 (GraphPad, San Diego, CA). Every dataset was tested for statistical normality and this information was used to apply the appropriate (parametric or nonparametric) statistical test. P values < 0.05 were considered significant; significance values are indicated as * p < 0.05, ** p < 0.01, *** p < 0.001, **** p < 0.0001.

## Results

### Generation and characterization of a 293T-Spike cell line

To develop a high-throughput flow cytometry assay able to detect anti-SARS-CoV-2 S antibodies in plasma from convalescent donors, we generated a cell line stably expressing the full-length S glycoprotein. Third-generation transgenic lentiviruses encoding for SARS-CoV-2 S were used to transduce 293T cells. Since the S glycoprotein is fused to a C-terminal GFP tag, 293T-Spike cells were sorted by flow cytometry based on GFP expression. The presence of cell-surface S was confirmed using the anti-RBD CR3022 monoclonal Ab and plasma from SARS-CoV-2 infected individuals. Specificity was confirmed using pre-pandemic healthy donor plasma (Figure 1A). For our high-throughput flow cytometry-based assay, parental 293T and 293T-Spike cells were mixed at an equal ratio and incubated with plasma from convalescent donors. Spike-specific antibodies were detected by adding a fluorescent anti-human IgG (H+L) secondary antibody. The signal was measured by flow cytometry and background signal measured on parental 293T cells (GFP negative) was subtracted for specificity. Signal obtained with plasma from 10 COVID-19 negative donors were used to define a limit of detection for seropositivity (Figure 1B, C).

**Figure 1.**
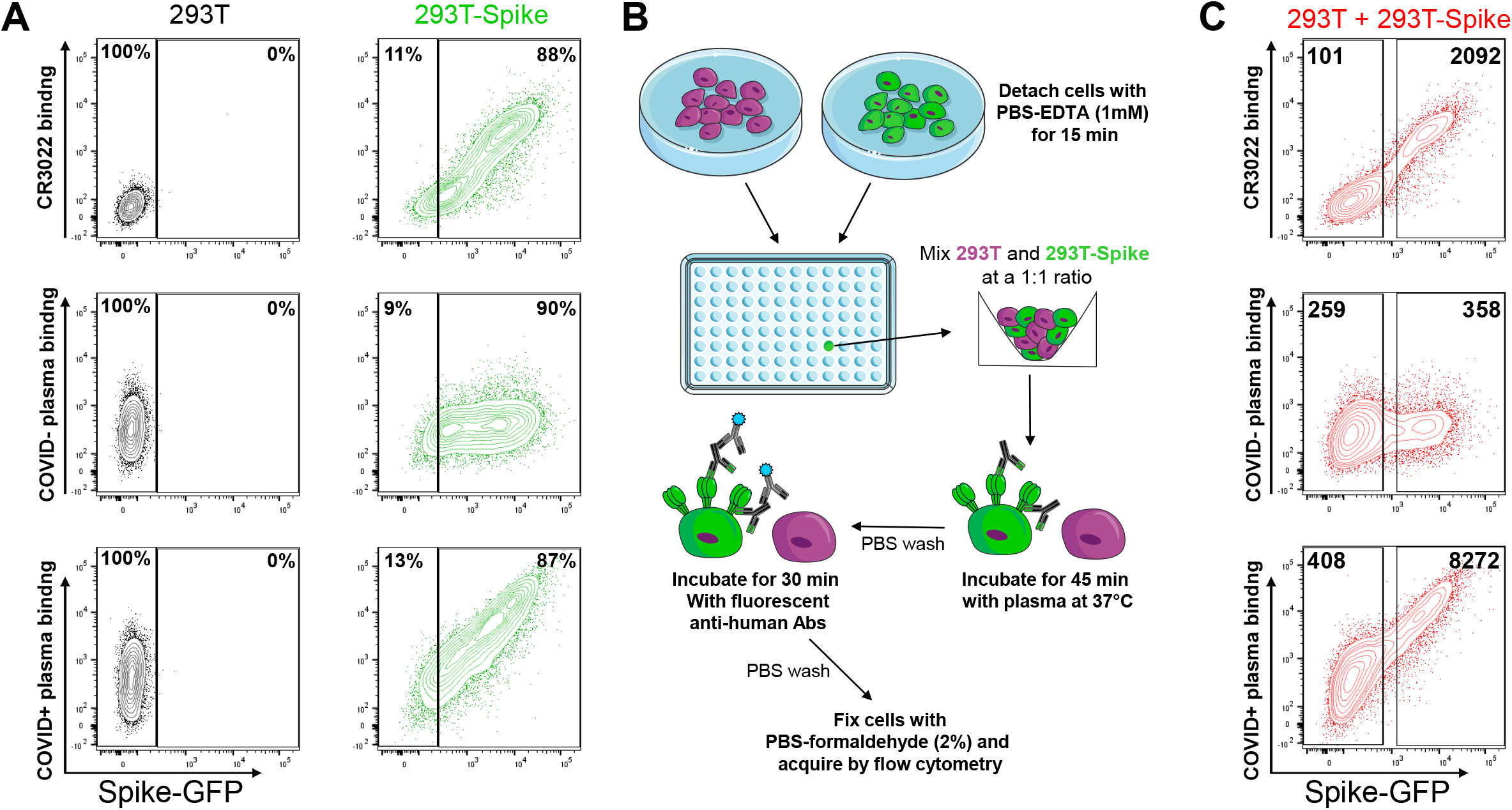
Characterization of the 293T-Spike cell line. (A) Dot plots depicting representative stainings of the parental 293T (left) or the 293T-Spike cell lines (right) using CR3022 mAb, a representative COVID-19 negative and COVID-19 positive plasma. Percentages represent the proportion of GFP+ and GFP-cells on the total cell population. (B) A schematic representation of the experimental procedures used to perform high-throughput screening (HTS) of plasma samples for their specific binding to SARS-CoV-2 Spike. (C) Dot plots depicting representative staining of pooled cell lines used for HTS assay (equal ratio of parental 293T (GFP-) and the 293T-Spike cells (GFP+)) using CR3022 mAb, a COVID-19 negative plasma and a COVID-19 positive plasma. Median fluorescence intensities (MFI) obtained on GFP- and GFP+ cell populations are indicated.

### Longitudinal decline of Spike-specific antibodies in plasma from convalescent donors

Recently, a longitudinal analysis was performed to measure the RBD-specific antibody response in convalescent plasma from 33 to 114 days post-symptom onset using a semi-quantitative ELISA ^18^. This cohort consisted of 11 males and 4 females (median age of 56 years old) and plasma was donated at least 4 times. A decrease in RBD-specific antibody titers between the first and last donations was observed for all 15 donors tested and this decline was shown to depend on time post-recovery but not on the number of donations. To extend this observation beyond the RBD domain, we used our high-throughput flow-cytometry based assay using the 293T-Spike cells to measure the persistence of antibodies targeting the full-length SARS-CoV-2 Spike in these convalescent plasma samples. Antibodies against S also decreased over time in these plasma samples, with the decrease being significant ~74 days post-symptom onset onwards (Figure 2A). This finding was corroborated using a previously characterized flow-cytometry method to quantify SARS-CoV-2 Spike-specific antibodies using 293T cells transiently transfected with a plasmid encoding the full-length Spike ^10,11,19–21^ (Figure 2B), and the MFI obtained from both these methods correlated significantly (r = 0.9207, p<0.0001) (Figure 2C). Results obtained with both flow cytometry assays, using transduced or transfected 293T cells, also positively correlated with the levels of RBD-specific antibodies as quantified by ELISA in the recently published study using the same cohort ^18^ (Figure 2C). Of note, the decline of total anti-Spike antibodies did not correlate with the number of donations (r = 0.1379, p = 0.6217) but rather correlated with the time elapsed between onset of symptoms and last donation (r = 0.5645, p = 0.0284) (Figure 2D).

**Figure 2.**
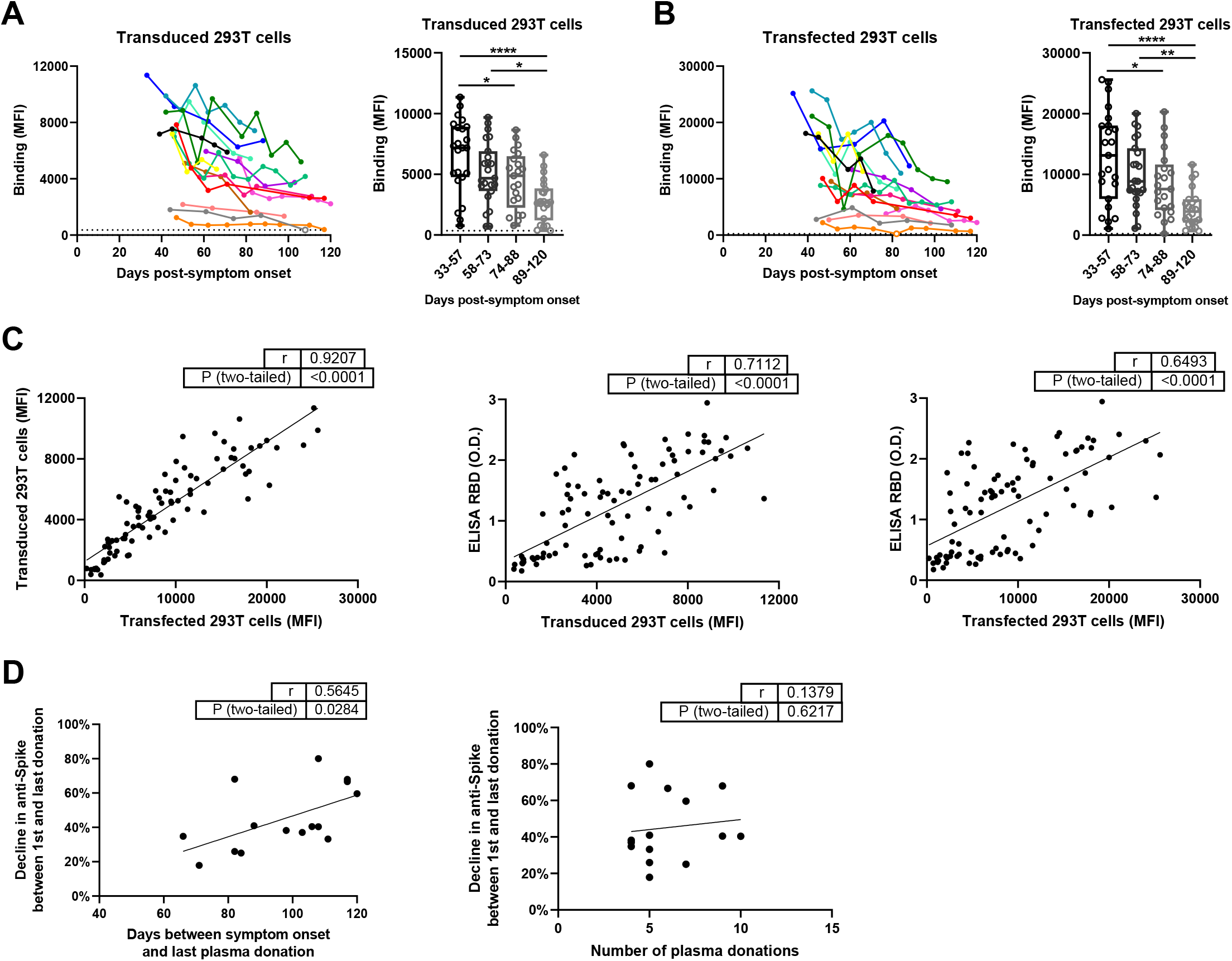
Decline of Spike-specific antibodies in longitudinal convalescent plasma. The level of anti-Spike antibodies in plasma from COVID+ donors was determined by flow cytometry using (A) 293T transduced cells or (B) 293T transfected cells expressing SARS-CoV-2 Spike. (A-B, left panels) Each curve represents the median fluorescence intensity (MFI) obtained with the plasma of one donor at every donation (4 to 10 donations per donor) as a function of the days after symptom onset. Undetectable measures are represented as white symbols, and limits of detection are plotted. (A-B, right panels) The time post-symptom onset (33-120 days) was divided in quartiles containing similar numbers (between 21 and 23) of plasma samples obtained from the 15 COVID-19 positive donors. Boxes and horizontal bars denote interquartile range (IQR) while horizontal line in boxes correspond to median of MFI values. Whisker endpoints are equal to the maximum and minimum values below or above the median ±1.5 times the IQR. Statistical significance was tested using one-way ANOVA with a Holm-Sidak post-test (* P < 0.05; ** P < 0.01; **** P < 0.0001. (C) Correlations between the levels of recognition of SARS-CoV-2 full-length Spike evaluated by flow cytometry using transduced or transfected 293T cells and levels of RBD recognition of SARS-CoV-2 RBD evaluated by indirect ELISA. (D) Correlations between the overall decline in Spike-specific antibody levels as measured by flow cytometry with transduced 293T cells (as calculated using the following formula: 1-[MFI at the last donation/ MFI obtained at first donation] x 100) and the number of days between symptom onset and the last donation or the number of donations by each donor. (C-D) Statistical significance was tested using a Pearson correlation test or a Spearman rank correlation test based on statistical normality.

## Discussion

There are many serodiagnosis platforms that have recently been approved for emergency use authorization (EUA) by the U.S Food and Drug Administration (FDA). In this study, we developed a high-throughput flow-cytometry based serodiagnosis tool by developing a cell line stably expressing the SARS-CoV-2 Spike to screen for anti-Spike antibodies in plasma of COVID-19 patients. Although our study shows data with plasma from only 15 donors, this assay can be readily adapted to a large-scale plasma screening with a high-throughput screening (HTS) plate reader for flow cytometry. In addition, we also expanded on recent findings showing a decrease in RBD-specific antibodies in convalescent plasma over time by showing that the level of antibodies targeting the full-length SARS-CoV-2 Spike also declines gradually after resolution of infection. These findings help to better understand the decline of humoral responses against the SARS-CoV-2 Spike and suggest that plasma should be collected rapidly after recovery from active infection in order to keep high levels of anti-Spike antibodies which are supposed to provide a clinical benefit in convalescent plasma transfer.

## Acknowledgements

The authors thank the CRCHUM Flow Cytometry Platform for technical assistance. The authors are grateful to the convalescent plasma donors who participated in this study and the Héma-Québec team involved in convalescent donor recruitment and plasma collection. We thank Dr. Stefan Pöhlmann (Georg-August University, Germany) for the plasmid coding SARS-CoV-2 S glycoproteins and Dr. M. Gordon Joyce (U.S. MHRP) for the monoclonal antibody CR3022. Figure 1B was prepared using images from Servier Medical Art by Servier, which is licensed under a Creative Commons Attribution 3.0 Unported License. This work was supported by le Ministère de I’Économie et de I’Innovation du Québec, Programme de soutien aux organismes de recherche et d’innovation to A.F. and by the Fondation du CHUM. This work was also supported by Canada’s COVID-19 Immunity Task Force (CITF), in collaboration with the Canadian Institutes of Health Research (CIHR) and a CIHR foundation grant #352417 to A.F. A.F. is the recipient of Canada Research Chair on Retroviral Entry no. RCHS0235 950-232424. S.P.A and J. Prévost are supported by CIHR fellowships. The funders had no role in study design, data collection and analysis, decision to publish, or preparation of the manuscript.

## Declaration of Interests

The authors declare no competing interests.

## Notes

### Competing Interest Statement

The authors have declared no competing interest.

